# Parallel evolution of two clades of a major Atlantic endemic *Vibrio parahaemolyticus* pathogen lineage by independent acquisition of related pathogenicity islands

**DOI:** 10.1101/155234

**Authors:** Feng Xu, Narjol Gonzalez-Escalona, Kevin P. Drees, Robert P. Sebra, Vaughn S. Cooper, Stephen H. Jones, Cheryl A. Whistler

**Author notes:** Current address: Microbiology and Molecular Genetics, University of Pittsburgh School of Medicine, Pittsburgh, PA.

## Abstract

Shellfish-transmitted *Vibrio parahaemolyticus* infections have recently increased from locations with historically low disease incidence, such as the Northeast United States (US). This change coincided with a bacterial population shift towards human pathogenic variants occurring in part through the introduction of several Pacific native lineages (ST36, ST43 and ST636) to near-shore areas off the Atlantic coast of the Northeast US. Concomitantly, ST631 emerged as a major endemic pathogen. Phylogenetic trees of clinical and environmental isolates indicated that two clades diverged from a common ST631 ancestor, and in each of these clades, a human pathogenic variant evolved independently through acquisition of distinct *Vibrio* pathogenicity islands (VPaI). These VPaI differ from each other and bear little resemblance to hemolysin-containing VPaI from isolates of the pandemic clonal complex. Clade I ST631 isolates either harbored no hemolysins, or contained a chromosome I-inserted island we call VPaIβ that encodes a type three secretion system (T3SS2β) typical of Trh hemolysin-producers. The more clinically prevalent and clonal ST631 clade II had an island we call VPaIγ that encodes both *tdh* and *trh* and that was inserted in chromosome II. VPaIγ was derived from VPaIβ but with some additional acquired elements in common with VPaI carried by pandemic isolates, exemplifying the mosaic nature of pathogenicity islands. Genomics comparisons and amplicon assays identified VPaIγ-type islands containing *tdh* inserted adjacent to the *ure* cluster in the three introduced Pacific and most other emergent lineages. that collectively cause 67% of Northeast US infections as of 2016.

**IMPORTANCE:** The availability of three different hemolysin genotypes in the ST631 lineage provided a unique opportunity to employ genome comparisons to further our understanding of the processes underlying pathogen evolution. The fact that two different pathogenic clades arose in parallel from the same potentially benign lineage by independent VPaI acquisition is surprising considering the historically low prevalence of community members harboring VPaI in waters along the Northeast US Coast that could serve as the source of this material. This illustrates a possible predisposition of some lineages to not only acquire foreign DNA but also to become human pathogens. Whereas the underlying cause for the expansion of *V. parahaemolyticus* lineages harboring VPaIγ along the US Atlantic coast and spread of this element to multiple lineages that underlies disease emergence is not known, this work underscores the need to define the environment factors that favor bacteria harboring VPaI in locations of emergent disease.

## Introduction

*Vibrio parahaemolyticus* is an emergent pathogen capable of causing human gastric infections when consumed, most often with contaminated shellfish (1, 2). Some human pathogenic *V. parahaemolyticus* variants evolve from diverse non-pathogenic communities through horizontal acquisition of *<u>V</u>ibrio* <u>pa</u>thogenicity <u>i</u>slands (VPaI) (3-5). Gastric pathogenic *V. parahaemolyticus* typically harbor islands with at least one of two types of horizontally acquired hemolysin genes *(tdh* and *trh)* that are routinely used for pathogen discrimination even though their role in disease appears modest (6-11). Most pathogenic *V. parahaemolyticus* isolates also carry accessory <u>t</u>ype <u>t</u>hree <u>s</u>ecretion <u>s</u>ystems (T3SS) that translocate effector proteins that contribute to host interaction (12-14). Two evolutionarily divergent horizontally-acquired accessory systems (T3SS2α or T3SS2β) contribute to human disease and are genetically linked to hemolysin genes (two *tdh* genes with T3SS2α, and *trh* with T3SS2β) in contiguous but distinct islands (4, 15-17). The first described *tdh*-harboring island [called by several different names including Vp-PAI (15), VPaI-7 (4), and *tdh*VPA (17)] from an Asian pandemic strain called RIMD 2210366 is fairly well-characterized (4, 5, 13, 18, 19). In contrast, islands containing T3SS2β linked to *trh* and a urease *(ure)* cluster, which confers a useful diagnostic phenotype, [where similar islands are described by others as Vp-PAI_TH3966_ (16), or *trh*VPA(17, 20)] have received only modest attention. Pathogenic variants harboring both *tdh* and *trh* are increasingly associated with disease in North America (21-26), and yet, to our knowledge, the exact configuration of hemolysin-associated VPaI(s) in isolates that contain both *tdh* and *trh* have not yet been described [although see (20)]. Thus it is unclear how virulence loci and islands in these emergent pathogen lineages carrying both hemolysins evolved and spread.

The expanding populations of *V. parahaemolyticus* have increased infections even in temperate regions previously only rarely impacted by this pathogen and where most environmental isolates harbor no known virulence determinants (27). A related complex of Asia-derived pandemic strains, most often identified as serotype O3:K6 and also known as sequence type (ST) 3 (based on allele combinations of seven housekeeping genes) causes the most disease globally (28). An unrelated Pacific native lineage called ST36 (also described as serotype O4:K12) currently dominates infections in North America, including from the Northeast United States (US) (21, 26, 29). The introduction of ST36 into the Atlantic Ocean by an unknown route precipitated a series of outbreaks from Atlantic shellfish starting in 2012 (29, 30). Prior to 2012, residential lineages contributed to low but increasing sporadic infection rates on the Northeast US coast (https://www.cdc.gov/vibrio/surveillance.html, 2017) (21), with ST631 emerging as the major lineage that is endemic to near-shore areas of the Atlantic Ocean bordering North America (the northwest Atlantic Ocean) (31). However, we previously identified a single ST631 isolate lacking hemolysins (21, 27) suggesting this pathogen lineage may have recently evolved through VPaI acquisition.

The goal of our study was to understand the genetic events and changing population context for the evolution of the ST631 pathogenic lineage. We conducted whole and core genome phylogenetic analysis of three environmental and 39 clinical ST631 isolates along with isolates from other emergent lineages from the region, which revealed two ST631 clades of common ancestry, from which human pathogens evolved in parallel. The single clade I clinical isolate acquired a *recA* gene insertion previously seen associated with Asian lineages, and had a VPaI that is typical of isolates harboring *trh* in the absence of *tdh.* In contrast, isolates from the clonal ST631 clade II that dominates Atlantic-derived ST631 infections (31) had a related but distinct VPaI. This VPaI contained a *tdh* gene and four associated hypothetical protein encoding genes inserted within, not next to, an existing ure-trh-T3SS2β island in close proximity to the *ure* cluster. Nearly all emergent resident and invasive lineages, including all three Pacific lineages (ST36, ST636 and ST43) contained islands that similarly had a *tdh* gene inserted within the VPaI in an identical location adjacent to the *ure* cluster providing a mechanism for simultaneous acquisition of both hemolysins with T3SS2β.

## RESULTS

### Atlantic endemic ST631 and several invasive lineages harboring both the *tdh* and *trh* hemolysin genes are clinically prevalent in four reporting Northeast US States

Ongoing analysis of clinical isolates revealed that even as the Pacific-derived ST36 lineage continued to dominate infections (50%), the endemic (autochthonous) ST631 lineage accounted for 14% of infections (Table 1). Concurrently, a limited number of other lineages contributed individually to fewer infections (≤3% each), among which were two lineages that have caused infections in the Pacific Northwest in prior decades: ST43 and ST636 (22, 23). ST43 and ST636 only recently (2013 and 2011 respectively) (21) have been linked to product harvested from waters along the Northeast US coast, and also caused infections in subsequent years. As is common among US clinical isolates, pathogenic isolates of all the aforementioned lineages harbor both the *tdh* and *trh* hemolysin genes (Table 1). Among environmental isolates, ST34 and ST674 are the most frequently recovered pathogen lineages but these caused comparatively few infections (Table 1). ST34 was first reported from the environment in 1998, from both the Gulf of Mexico and near-shore areas of MA, and was also recovered in NH in 2012 (21) suggesting it is an established resident in the region. ST674 which was first reported from an infection in Virginia in 2007 (32) was first recovered from the local environment in 2012 (www.pubmlst.org/vparahaemolyticus) (21). Notably even though all four ST674 environmental isolates, like ST34, harbored both hemolysin genes, the single ST674 clinical isolate (MAVP-21) lacked hemolysins (Table 1) (21). The decrease in clinical prevalence of *trh-*harboring Atlantic endemic ST1127, which caused no infections in the last three years, coincided with the increase in clinical prevalence of all three Pacific-derived lineages which harbor both hemolysins. Notably, very few other clinical isolates harbored *trh* in the absence of *tdh* and clinical isolates containing only *tdh* (i.e. ST1725) were extremely rare (Table 1). Concurrent with this shift in composition of clinical lineages that includes multiple Pacific-derived lineages, hemolysin producers have increased in relative abundance in nearshore areas of the region, where historically these represented ~1% of all isolates (27). Since 2012, hemolysin producers have been recovered more frequently, and in the last two years their proportion has increased by up to an order of magnitude (comprising as much as 10%) in some regional shellfish associated populations (data not shown).

**Table 1:** Clinical and environmental prevalence of emergent Northeast US *V. parahaemolyticus* lineages with associated virulence features.

| Sequence type<sup>a</sup> | Northeast US States<sup>b</sup> |  | MLST Database<sup>c</sup> |  | Hemolysin genotype | VPaI type<sup>d</sup> |
| --- | --- | --- | --- | --- | --- | --- |
|  | Clinical | Environmental | Clinical | Environmental |  |  |
| 3 | 2 | 0 | 217 | 33 | *tdh*<sup>+</sup> | $\alpha$ |
| 36 | 91 | 1 | 58 | 5 | *tdh*<sup>+</sup>*trh*<sup>+</sup> | $\gamma$ |
| 631 | 24 | 0 | 12 | 0 | *tdh*<sup>+</sup>*trh*<sup>+</sup> | $\gamma$ |
| | 1<sup>e</sup> | 2 | 0 | 0 | *trh*<sup>+</sup> | $\beta$ |
|  | 0 | 1 | 0 | 0 | neither | absent |
| 43 | 5 | 0 | 17 | 4 | *tdh*<sup>+</sup>*trh*<sup>+</sup> | $\gamma$ |
| 636 | 4 | 0 | 2 | 0 | *tdh*<sup>+</sup>*trh*<sup>+</sup> | $\gamma$ |
| 1127 | 4 | 0 | 0 | 0 | *trh*<sup>+</sup> | $\beta$ |
| 110 | 3 | 0 | 0 | 1 | *tdh*<sup>+</sup>*trh*<sup>+</sup> | $\gamma$ |
| 34/324 | 2 | 2 | 4 | 19 | *tdh*<sup>+</sup>*trh*<sup>+</sup> | $\gamma$ |
| 674 | 0 | 4 | 1 | 20 | *tdh*<sup>+</sup>*trh*<sup>+</sup> | $\gamma$ |
|  | 1 | 0 | 0 | 0 | neither | absent |
| 308 | 2 | 0 | 0 | 2 | *tdh*<sup>+</sup>*trh*<sup>+</sup> | $\gamma$ |
| 12 | 2 | 0 | 0 | 4 | *trh*<sup>+</sup> | $\beta$ |
| 162 | 2 | 0 | 1 | 1 | neither | absent |
| 194 | 2 | 0 | 1 | 0 | neither | absent |
| 809 | 2 | 0 | 0 | 1 | *trh*<sup>+</sup> | $\beta$ |
| 1716 | 2 | 0 | 0 | 0 | *trh*<sup>+</sup> | $\beta$ |
| 1123 | 1 | 1 | 0 | 0 | *trh*<sup>+</sup> | $\beta$ |
| 8 | 1 | 0 | 13 | 5 | *trh*<sup>+</sup> | $\beta$ |
| 23 | 1 | 0 | 0 | 3 | *tdh*<sup>+</sup>*trh*<sup>+</sup> | $\gamma$ |
| 749 | 1 | 0 | 1 | 0 | *tdh*<sup>+</sup>*trh*<sup>+</sup> | $\gamma$ |
| 1295 | 1 | 0 | 0 | 1 | neither | absent |
| 134 | 1 | 0 | 1 | 0 | neither | absent |
| 741 | 1 | 0 | 0 | 1 | neither | absent |
| 98 | 1 | 0 | 0 | 1 | *trh*<sup>+</sup> | $\beta$ |
| 1205 | 1 | 0 | 0 | 1 | neither | absent |
| 1561 | 1 | 0 | 0 | 0 | neither | absent |
| 1717 | 1 | 0 | 0 | 0 | neither | absent |
| 1725 | 1 | 0 | 0 | 0 | *tdh*<sup>+</sup> | $\alpha$ |

<sup>a</sup> Some clinical isolates had insufficient sequencing coverage to determine sequence type and included eight *tdh*<sup>+</sup>*trh*<sup>+</sup> isolates, one *tdh*<sup>+</sup> isolate, four *trh*<sup>+</sup> isolates, and 11 isolates without hemolysins, some of which were from wound infections. Two wound infection isolates lacking hemolysins were of known sequence types and are not listed above.
<sup>b</sup> Data generated from all available gastric infection clinical and environmental isolates four reporting Northeast US States including ME, NH, MA, and CT between 2010 and 2016.
<sup>c</sup> http://pubmlst.org/vparahaemolyticus, 2017 (36, 58)
<sup>d</sup> Presence of the VPαIγ architecture was determined by PacBio genome sequencing of isolate MAVP-Q and MAVP-26, whereas for other isolates, identification of VPαI type was determined through illumina genome sequencing, PCR amplification and Sanger sequencing.
<sup>e</sup> This single isolate harbors a *recA* allele (allele 21) typical of ST631 fused to allele 107 through an insertion event, generating a hybrid allele previously described (33).

### A single clinical ST631 lineage isolate with an unusual *recA* allele harbors *trh* in the absence of *tdh*

Employing ST631-specific marker-based assays (see methods), we identified two additional 2015 environmental isolates (one from NH and one from MA) and one additional 2011 local-source clinical isolate (MAVP-R) (21) with a hemolysin profile *(trh^+^* without *tdh)* that is atypical of the ST631 lineage (Table 1). Although analysis of the seven-housekeeping gene allele combination confirmed the environmental isolates were indeed ST631, MAVP-R was not ST631 based on only one locus: *recA.* Examination of the *recA* locus of MAVP-R uncovered a large insertion within the ancestral ST631 *recA* gene (allele recA21; www.pubmlst.org/vparahemolyticus) incorporating an intact but different *recA* gene into the locus [allele recA107(33)] and fragmenting the ancestral gene (Fig. 1). The insertion in the ancestral *recA* gene in MAVP-R is identical to one observed in the *recA* locus of two Hong Kong isolates (isolates S130 and S134) and similar to the one in isolate 090-96 (ST189a) isolated in Peru but believed to have originated in Asia (33).

**Figure 1.**
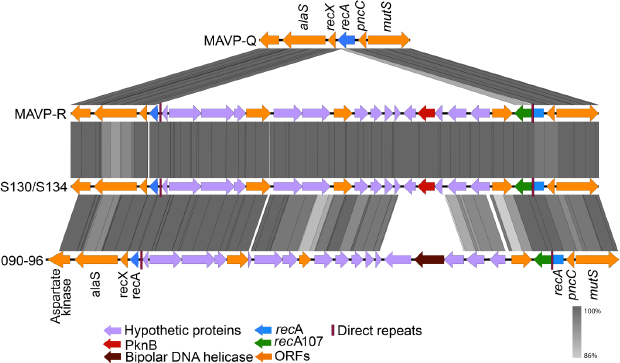
Schematic of a horizontally acquired insertion in the recA-encoding region of MAVP-R. Sequences of the *recA* gene and flanking region from MAVP-Q (reference ST631 genome), MAVP-R, Asia-derived isolates S130/S134 and Peru-derived isolate 090-96 were extracted and aligned. Open reading frames designated with arrows and illustrated by representative colors to highlight homologous and unique genes. The *%* similarity between homologs is illustrated by grey bars.

### ST631 forms two divergent clades

The existence of three different hemolysin profiles (Table 1) among all available ST631 draft genomes suggested there could be more than one ST631 lineage. Therefore we evaluated whole genome maximum likelihood (ML) phylogenies of select ST631 isolates and all other lineages causing two or more infections reported in four Northeast US States to evaluate whether there was more than one ST631 lineage (Table 1) (Fig. 2). The phylogenetic tree showed that ST631 isolates, regardless of their hemolysin genotype, clustered together but they formed two distinct clades, indicative of common ancestry (Fig. 2). Clade I harbored either *trh* or no hemolysins and consisted of all three environmental isolates which were from MA and NH, and the single clinical isolate MAVP-R, whereas clade II consisted of all other isolates all of which harbor both hemolysins. The two distinct ST631 clades shared 85% of their DNA in common and displayed polymorphisms in ≤12% of the shared DNA content. The most closely related sister lineage to ST631 was formed by *trh*-harboring ST1127 isolates that have been exclusively reported from clinical sources in the Northeast US (21).

**Figure 2.**
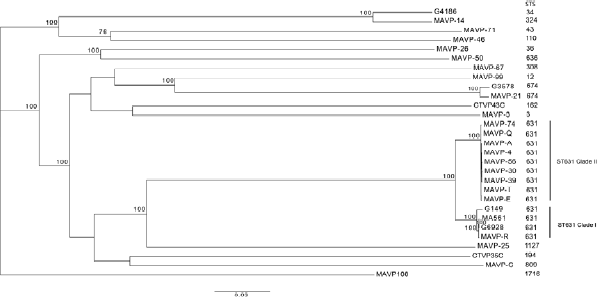
Phylogenetic relationships of *V. parahaemolyticus* lineages and identification of distinct ST631 clades. An ML phylogeny of representative *V. parahaemolyticus* genomes of clinical isolates causing two or more infections was built on whole genome SNPs identified by reference-free comparisons as described in the methods. The branch length represents the number of nucleotide substitutions per site. Numbers at nodes represent percent bootstrap support where unlabeled nodes had bootstraps of less than 70.

We next evaluated the relationships of all available ST631 isolate genomes at NCBI and sequenced by us (Supplemental Table 1) using a custom core genome multi-locus sequence typing (cgMLST) method as previously described (31). Minimum spanning trees built from core genome loci from 42 ST631 isolates indicated that only 390 loci varied between the most closely related isolate of clade I (MAVP-L) and clade II (G6928) (Fig. 3). The most distantly related isolates within clade I (G149 and MAVP-R) exhibited 80 core genome loci differences whereas clade II is clonal with only 51 variant loci between the most divergent isolates: clinical isolate 09-4436 and environmental isolate S487-4, both reported from PEI Canada (Fig. 3) (31).

**Figure 3.**
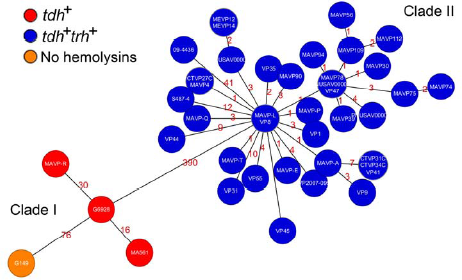
Minimum spanning tree relationships among clade I and clade II ST631. A cgMLST core gene-by-gene analysis (excluding accessory genes) was performed and SNPs were identified within different alleles. The numbers above the connected lines (not to scale) represent SNP differences. The isolates are colored based on different hemolysin genotypes as labeled.

### Each ST631 clade independently acquired a distinct pathogenicity island positioned on different chromosomes

Given the variation in ST631, comparisons between these isolates could elucidate the events that led not only to the evolution of two pathogenic clades but also address unresolved questions about the unique configurations and contents of pathogenicity islands in western Atlantic Ocean emergent lineages. The physical proximity of *tdh* with the *ure* cluster and *trh*, and the co-occurrence of *tdh* with T3SS2β reported in many *tdh^+^/trh^+^* clinical isolates suggested *tdh* could be harbored within or next to the same pathogenicity island harboring *trh* in at least some lineages as was previously suggested (20, 24, 34).

To identify the location and determine the architecture of the pathogenicity elements harboring hemolysin genes, we generated high quality annotated genomes for the clade I ST631 isolate MAVP-R and clade II ST631 isolate MAVP-Q (both reported in 2011 from MA) employing PacBio sequencing. The pathogenicity island regions in these isolates genomes were extracted, aligned, and the contents compared with pathogenicity island harboring two *tdh* genes [previously called Vp-PAI (15), VPaI-7 (4) and *tdh*VPA(17)] from RIMD 2210366 and Vp-PAI_TH3996_ (16) [also called *trh*VPI (17)] harboring *trh* (Supplemental Table 2). This comparison revealed that MAVP-R harbored a pathogenicity island typical of *trh*-containing isolates that includes a linked *ure* cluster and T3SS2β that is orthologous, with the exception of few unique regions, with Vp-PAI_TH3996_ (16) (Supplemental Table 2 and Fig. 4). Because the lack of convention in uniformly naming syntenous islands that distinguish them from distinctive and yet functionally analogous islands can impede communication, we hereafter will consistently reference the same island by a common descriptive name regardless of isolate lineage. Hereafter we will refer to islands sharing the same general configuration to that in MAVP-R by the name VPaIβ, and refer to *tdh*-containing islands similar to that described in strain RIMD 2210366 by the name VPaIa, regardless of bacterial isolate background. We adopted this simplified nomenclature in reference to the version of the key virulence determinant carried in the islands (T3SS2α and T3SS2β) in the two already described island types. This scheme importantly accommodates naming of additional uniquely-configured islands as they are identified. As noted previously (16, 17, 20), VPalβ is dissimilar to VPala in most gene content with ~ 78 ORFs unique to VPaIβ (where the number of identified ORFs used for comparison can differ slightly depending on which annotation program is applied) (Supplemental Table 2, Fig. 4). Even so, VPalβ had many homologous genes of varying sequence identity (n=~38 ORFs, excluding *tdh* homology with *trh)* when compared to VPalα (Supplemental Table 2, Fig. 4)(4, 5, 16). Identification of some homologs required that we relax matching to 50% such as for the divergent, but homologous T3SS2α and T3SS2β genes encoding the apparatus, chaperones, and some shared effectors (Supplemental Table 2). No homolog of the T3SS2α effector gene *vopZ* was identified, but a single ORF whose deduced protein sequence bears only 27% identity with VopZ is located in its place (Fig. 2 and Supplemental Table 2). VPaIβ from strain TH3996 and VPaIα from pandemic strain RIMD 2210633 are inserted in an identical location in chromosome II adjacent to an Acyl-CoA hydrolase-encoding gene. In contrast the VPaIβs in MAVP-R, ST1127 isolate MAVP-25, and Asia-derived AQ4037 are in chromosome I, in each case in the same insertion location identified for strain AQ4037 (17).

**Figure 4.**
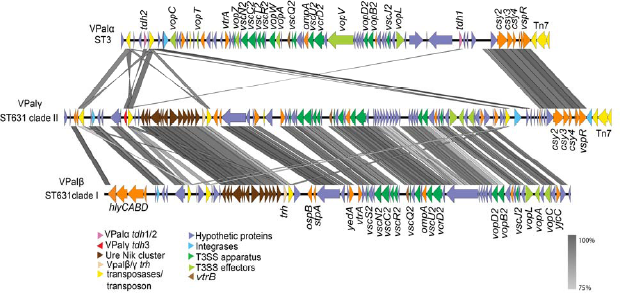
Comparisons of the pathogenicity islands containing hemolysins and T3SS2. Sequences of VPal were extracted from select genomes and aligned. VPala was derived from ST3 strain RIMD2210633, VPaly was derived from ST631 clade II isolate MAVP-Q, and VPaIβ was derived from ST631 clade I isolate MAVP-R. ORFs are depicted in defined colors and similarities (≥75%) among ORFs are illustrated in grey blocks. Homologs between VPalα and VPalβ/γ (50Ↄ75% identity) are named and listed in Supplemental Table 2.

MAVP-Q contained both *tdh* and *trh* within the same contiguous unique VPaI (hereafter called VPaIy) that shared features with both VPaIα and VPaIβ (Fig. 4, Supplemental Table 2). Specifically, VPaIγ had a core that with few exceptions was orthologous in content and syntenous with VPaIβ from MAVP-R (Fig. 4) with only a few exceptions. VPaIγ displays high conservation with VPaIa near its 3' end, as has been described in other draft *tdh^+^trh^+^* harboring genomes (20) as well as in the VPaIβ island of strain TH3996, although the presence of this element may not be typical of VPaIβ (e.g. it is absent in the islands from AQ4037 (17), MAVP-R and MAVP-25). The VPaIγ also contained a *tdh* gene homologous to *tdh2* (also called *tdhA)* from VPaIα (98.6%) near its 5' end but not at the 5' terminus of the island (Fig. 4). Rather, the DNA flanking both sides of the *tdh* gene in VPalγ was conserved in VPalβ of MAVP-R and absent from VPaIa, (Fig. 4). Analysis of 300 genomes of *V. parahaemolyticus* (representing a minimum of 28 distinct sequence types) of sufficient quality for analysis confirmed that the module of four hypothetical proteins preceding the *tdh2* homolog was present only in *trh-* harboring genomes, but not in genomes harboring *tdh* in the absence of *trh* (i.e. VPaIα containing genomes), providing evidence that the *tdh* gene was acquired horizontally by insertion into, not next to, an existing VPaIβ, perhaps through activity of the adjacent transposase gene (11) (Supplemental Table 3, Supplemental fig. 1, and data not shown). Like with VPaIa from RIMD 2210633, and VPaIβ of TH3996, VPaIγ of clade II ST631 is located in a conserved location of chromosome II, adjacent to an Acyl-CoA hydrolase-encoding gene.

The final environmental ST631 clade I isolate that lacked hemolysins, G149, had no VPaIα, β or γ elements in its genome. Close examination of the DNA corresponding to the VPaI insertion sites in either chromosome revealed no remnants of these islands in either chromosomal location indicating this isolate likely never acquired a pathogenicity island (Supplemental Fig 2 and data not shown). Because clade I isolate G149 lacked these islands, this could be the ancestral state of the ST631 lineage (21).

### Most clinically prevalent isolates from the Northeast US harbor similar contiguous pathogenicity islands containing *tdh* inserted in the same location of their VPaI

We next asked which isolates from other lineages likely residing within the mixed population with ST631 in near-shore areas of the Northeast US harbored islands of similar structure to VPaIγ that contain both hemolysin genes. Assembly of short-read sequences into contigs that cover the full length of VPaI which is necessary for comparative analysis of entire island configuration was impeded by the fact that homologous transposase sequences and other sequences were repeated multiple times throughout the island. Therefore, we determine whether other lineages harboring both hemolysin genes harbor *tdh* in the same island location, between the conserved VPalβ/γ module of four hypothetical proteins (to the left or 5' of *tdh)* and the *ure* cluster (to the right or 3' of *tdh)* (Fig. 4) by combining bioinformatics analysis of sequenced genomes with amplicon assays (Supplemental Fig 1). First we analyzed assembled draft genomes for *tdh* co-occurrence and proximity with the four adjacent hypothetical protein encoding genes that are absent in VPalβ but present in VPaly (See Methods). Every emergent pathogenic lineage of the Northeast US (Table 1) harboring both *tdh* and *trh* carried homologous DNA corresponding to all four hypothetical proteins adjacent to the *tdh* gene in a contiguous segment (Supplemental Table 3). To determine whether *tdh* was also adjacent to the *ure* cluster in these same isolates we next designed specific flanking primers and amplified the unique juncture between the tdh-containing transposon associated module and the *ure* cluster for all clinical isolates harboring both *tdh* and *trh* (See Methods) (Supplemental Fig 1). The results were congruent with our bioinformatics assessment (Supplemental Table 3), and demonstrated that isolates from all emergent pathogenic lineages harboring both hemolysins have *tdh* inserted in close proximity to an *ure* cluster in a configuration similar to VPalγ from MAVP-Q (Fig. 5, Table 1). This confirmed that these isolates harboring both hemolysins harbor *tdh* within, and not next to, the same VPal thereby facilitating simultaneous acquisition of both hemolysin genes.

**Figure 5.**
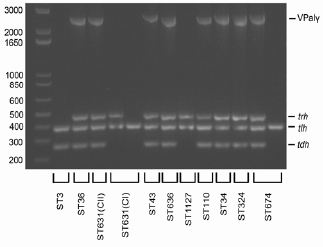
Distribution of VPaIγ in emergent pathogen lineages. The presence of *tdh, trh* and VPaIy along with positive control *tlh* was determined by PCR amplification using gene-specific primers and visualized on a 1.2% agarose gel. The order from left to right is 1kb+ ladder, ST3 (MAVP-C), ST36 (MAVP-26), ST631 CII (clade II isolate MAVP-Q), ST631 CI (clade I isolates MAVP-R and G149), ST43 (MAVP-71), ST636 (MAVP-50), ST1127 (MAVP-M), ST110 (MAVP-46), ST34 (CTVP19C), ST324 (MAVP-14), and ST674 (CT4291, MAVP21). The corresponding sizes of the ladder fragments are as labeled to the left and the identity of the amplicons listed to the right of the gel image.

## DISCUSSION

Even preceding the increased illnesses from Pacific-invasive lineages, two different clades of the predominant endemic Atlantic lineage of pathogenic *V. parahaemolyticus,* ST631 (31) evolved and contributed to a rise in sporadic illnesses in the four reporting Northeast US States (Table 1, Fig. 2 & 3). Several lines of evidence support the interpretation of parallel pathogen evolution. The two lineages exhibit differences in both clinical and environmental prevalence suggesting the pathogenic variants of each clade have not evolved the same degree of virulence (Table 1). Pathogenic members in each lineage also acquired different pathogenicity islands with different hemolysin gene content (Fig. 2 & 3). Although it was a formal possibility ^1^ that ST631 clade II evolved from clade I by independent horizontal acquisition of *tdh* into its existing VPaIβ, it is notable that other resident and even invasive lineages now in the Atlantic harbor VPaIγ with *tdh* and four additional co-occurring ORFs inserted into the same location of the island, suggesting a common evolutionary origin of this hybrid-type island (Fig. 4 and Supplemental Fig. 1). Finally, each of the two clades harbor VPaI insertions on different chromosomes: the less clinically prevalent ST631 clade I contains three isolates that harbor i VPaIβ in chromosome I (Fig. 3) and a single environmental isolate lacking any island (Table 1, supplemental Fig. 2), whereas the clonal ST631 clade II isolates all harbor VPaIγ on chromosome II.

Given that several other resident lineages harbor similar β and γ-type VPaI, pathogens in each clade could have acquired their islands from the reservoir of resident bacteria already i circulating in the Atlantic even before the presume arrival of invasive Pacific lineages. Several well-documented members of the Gulf of Mexico *V. parahaemolyticus* population (35-37) may also have expanded their range through movement of ocean currents and could be the source for these VPaI (Table 1, Fig. 5). But historically, hemolysin producers were extremely rare in near shore areas of the Atlantic US coast (25) and represented only about ~1% of isolates in an i estuary of NH as of a decade ago (27) limiting the potential for interacting partners or sources for acquired VPaI. Given this historical context, it is remarkable that two different clades from the same lineage independently acquired different VPaI-which for clade II ST631 occurred prior to 2007 ‐well before the recent shift in abundance of hemolysin producers.

The parallel evolution of two different lineages through lateral DNA acquisition alludes to the possibility that as-yet-undefined attributes may increase the chances of acquisition or prime some bacterial lineages (such as ST631) to more readily acquire and maintain genetic material or become pathogenic upon island acquisition. Even though the ecological niche in which horizontal island acquisition took place is unknown, it is conceivable that co-colonization of hosts or substrates favorable to the growth of ST631 and hemolysin producers may have facilitated island movement. Certainly, association of bacteria with specific marine substrates such as chitinous surfaces of plankton that also induce a natural state of competence could promote lateral transfer through close contact between the progenitors of the pathogenic subpopulation of each clade and island donors (3, 38, 39). Alternatively, conjugative plasmids or transducing phage could have been the agents of island delivery. The finding that the only clinical clade I isolate, MAVP-R, also harbors a second horizontal insertion in its *recA* locus that matched one previously found in Asia-derived strains (33) indicates it acquired more than one segment of foreign DNA during its evolution as a pathogen (Fig. 1) further illustrating that mechanisms that facilitate DNA transfer and acquisition may both have been at play. It also suggests that horizontal transfer of DNA from introduced bacteria not yet detected in the Atlantic could add to the genetic material available for pathogen evolution from Atlantic Ocean populations. The more detailed molecular epidemiological, comparative genomics, and functional analyses necessary to assess the impact of introduced pathogens on resident Atlantic lineages are warranted given this evidence and the documented introduction of multiple Pacific-derived lineages in the region (Table 1).

There has been some consideration of the roles of human virulence determinants in ecological fitness, but the natural context of pathogenic *V. parahaemolyticus* evolution is still unknown (40-42). Whereas *tdh* and T3SS2α each may promote growth when bacteria are under predation, isolates that carry *trh*-containing islands (which likely also have T3SS2β) do not derive similar benefits from their islands (43). This is surprising considering the islands encode several homologous effectors (Fig. 4 and Supplemental Table 2) that don't have an established role in enteric disease but they could alternatively or additionally mediate eukaryotic cell interactions with natural hosts thereby promoting environmental fitness (13, 14). But these islands also lack homologous open for the VPalα effector that is most closely associated with enteric disease: *vopZ* (11) (Fig. 4 and Supplemental Table 2). The general lack of knowledge of unique T3SS2β effectors and other gene function in these islands (Fig. 4 and Supplemental Table 2) even with regard to enteric disease, limits comparative analysis with the well-studied and functionally defined VPalα which could elucidate the bases for pathogen evolution. The higher clinical prevalence of clade II ST631 than clade I which has also been recovered on more than one occasion from the environment (Table 1) could indicate that VPaly confers greater virulence potential than VPaIβ, perhaps owing to the presence of *tdh,* a known virulence factor (1, 7, 44). However, the resident community members in both the Pacific and the Atlantic Ocean that harbor *tdh* and T3SS2α comparatively rarely cause human infections (21-23). The unique environmental conditions that underlie pathogen success from northern latitudes that favors bacteria with VPaIβ and VPaIγ including two different ST631 lineages suggests the shared content of these islands could confer abilities that are distinct from VPaIa which could underlie the repeated acquisition and maintenance of these related islands by so many different lineages now present in near-shore areas of the Northeast US.

## MATERIALS AND METHODS

### Bacteria isolates, media and growth conditions

*V. parahaemolyticus* clinical isolates for this study were provided by cooperating public health laboratories in Massachusetts, New Hampshire, Maine, and Connecticut whereas a select number of environmental isolates were enriched from estuarine substrates as described (21). Detailed information about these isolates was described previously (31) and listed in Supplemental Table 1. Isolates were routinely cultured in Heart Infusion (HI) media supplemented with NaCl at 37°C as described (21).

### Whole genome sequencing, assembly, annotation and sequence type identification

Genomic DNA was extracted using the Wizard Genomic DNA purification Kit (Promega, Madison WI USA) or by organic extraction (21). The quality genomic DNA was determined by spectrophotometric measurements by NanoDrop (ThermalFisher, Waltham MA USA). Libraries for DNA sequencing were prepared using a high-throughput Nextera DNA preparation protocol (45) using an optimal DNA concentration of 2ng/μl. Genomic DNA was sequenced using an Illumina – HiSeq2500 device at the Hubbard Center for Genome Studies at the University of New Hampshire, using a 150bp paired-end library. *De novo* assembly was performed using the A5 pipeline (46), and the assemblies annotated with Prokka1.9 using the "genus" option and selecting *"Vibrio”* for the reference database (47). The sequence types were subsequently determined using the SRST2 pipeline (48). The sequence type of each genome was determined when using *V. parahaemolyticus* as the database (https://pubmlst.org/vparahaemolyticus/). For most isolates where the combination of each allele was not found in the database representing novel sequence types, the genome was submitted for a new sequence type designation (www.pubmlst.org/vparahaemolyticus).

Isolates MAVP-Q and MAVP-R were sequenced using the Pacific Biosciences RSII technology. Using between 3.7-5.3 μg DNA, the library preparation and sequencing was performed according to the manufacturer’s instructions (Pacific Biosciences, Menlo Park CA, USA) and reflects the P5-C3 sequencing enzyme and chemistry for MAVP-Q isolate and the P6-C4 configuration for MAVP-R. The mass of double-stranded DNA was determined by Qubit (Waltham, MA USA) and the sample diluted to a final concentration of 33 μg / μL in a volume of 150 μL elution buffer (Qiagen, Germantown MD USA). The DNA was sheared for 60 seconds at 4500 rpm in a G-tube spin column (Covaris, Wobrun MA USA) which was subsequently flipped and re-spun for another 60 seconds at 4500 rpm resulting in a ~20,000 bp DNA verified using a DNA 12000 Bioanalyzer gel chip (Agilent, Santa Clara, CA USA). The sheared DNA isolate was then re-purified using a 0.45X AMPure XP purification step (Beckman Coulter, Indianapolis IN USA). The DNA was repaired by incubation in DNA Damage Repair solution. The library was again purified using 0.45X Ampure XP and SMRTbell adapters ligated to the ends of the DNA at 25°C overnight. The library was treated with an exonuclease cocktail (1.81 U/μL Exo III 18 and 0.18 U/μL Exo VII) at 37°C for 1 hour to remove un-ligated DNA fragments. Two additional 0.45X Ampure XP purifications steps were performed to remove <2000 bp molecular weight DNA and organic contaminant.

Upon completion of library construction, samples were validated using an Agilent DNA 12000 gel chip. The isolate library was subjected to additional size selection to the range of 7,000 bp – 50,000 bp to remove any SMRTbells < 5,000 bp using Sage Science Blue Pippin 0.75% agarose cassettes to maximize the SMRTbell sub-read length for optimal *de novo* assembly. Size-selection was confirmed by Bio-Analysis and the mass was quantified using the Qubit assay. Primer was then annealed to the library (80°C for 2 minute 30 followed by decreasing the temperature by 0.1°/s to 25°C). The polymerase-template complex was then bound to the P5 or P6 enzyme using a ratio of 10:1 polymerase to SMRTbell at 0.5 nM for 4 hours at 30°C and then held at 4°C until ready for magbead loading, prior to sequencing. The magnetic bead-loading step was conducted at 4°C for 60-minutes per manufacturer’s guidelines. The magbead-loaded, polymerase-bound, SMRTbell libraries were placed onto the RSII machine at a sequencing concentration of 110-150 pM and configured for a 180-minute continuous sequencing run. Long read assemblies were constructed using HGAP version 2.3.0 for *de novo* assembly generation. Further, hybrid assemblies were generated and error corrected with illumina raw reads using Pilon v1.20 (49).

### Lineage-specific marker-based assays

To more rapidly identify ST631 isolates from clinical and environmental collections we developed PCR-amplicon assays to unique gene content in ST631. Whole genome comparisons were performed on MAVP-Q (a ST631 clinical isolate), G149 (a ST631 environmental isolate), MAVP-26 (ST36), RIMD2210633 (ST3), and AQ4037 (ST96) (Supplemental Fig. 3). A total of 26 distinct genomic regions, each greater than 1kb in size, were present in MAVP-Q but absent in other comparator genomes, including environmental ST631 that lacks hemolysins (G149) (Supplemental Fig. 3). Within a large genomic island ~37.6 Kb in length with an integrase at one terminus and an overall lower GC content (40.6% compared to 45.8% for the genome) a single ORF homologous to restriction endonucleases (AB831_06355) that was restricted to clinical ST631 isolates in our collection and publicly available draft genomes (n=693) (http://www.ncbi.nlm.nih.gov/genome/691, 2017) was selected as a suitable amplicon target. The distribution of this locus was further analyzed using the BLAST algorithm by a query against the nucleotide collection, the non-redundant protein sequences, and against the genus *Vibrio* (taxid: 662), excluding *V. parahaemolyticus* (taxid: 691), using the default settings for BLASTn (50). Similar approaches were applied to identify ST631 diagnostic loci inclusive of the single environmental isolate (G149), which identified a hypothetical protein encoding region (AB831_06535) (ST631env). Oligonucleotide primers were designed to amplify the diagnostic regions including AB831_06355 using primers *ST631end* F (5'AGTTCATCAGGTAGAGAGTTAGAGGA3') and ST631endR (5’TCTTCGTTACCATAGTATGAGCCA3’) which produces and amplicon of c.a. 494bp, and AB831_06535 using primers ST631envF (5'TGGGCGTTAGGCTTTGC3') and ST631-envR (5'GGGCTTCTACGACTTTCTGCT3') producing an amplicon of 497bp.

Amplification of diagnostic loci was evaluated in individual assays using genomic DNA from positive and negative controls: MAVP-Q and G149 (ST631), G4186 (ST34), G3578 (ST674), and MAVP-M (ST1127), MAVP-26 (ST36) and G61 (ST1125). Amplification of specific sequence types were performed with Accustart enzyme mix on purified DNA. Cycling was performed with an initial denaturation at 94°C for 3 min., followed by 30 cycles of a denaturation at 94°C for 1 min, annealing at 55°C for 1 min, and amplification at 72°C for 30s with a final elongation at 72°C for 5 min. The primer pairs only produced amplicons from template DNA from ST631 and each was the expected size (data not shown, and Supplemental Fig. 3). Amplicon assays were applied to 208 clinical isolates from the Northeast US States (ME, NH, MA and CT) and 1140 environmental isolates collected from 2015-2016 from NH and MA. These assays identified all known ST631 clinical isolates with 100% specificity and also identified an additional 7 *tdh^+^trh^+^* clinical isolates (ST631*end* and ST631env positive), and two environmental (ST631end negative and ST631env positive) isolates from our archived collection. Each, with the exception of MAVP-R, was subsequently confirmed to be ST631 by seven-locus MLST (www.pubmlst.org).

### Examination of *recA* allele and adjacent sequences

The PacBio sequenced genome of MAVP-R, contig 000001 (Accession No. MPPP00000000) that contained the *recA* gene, was annotated using PROKKA1.9 (47). The sequences of *recA* and its surrounding DNA was then compared to the contig containing *recA* region from isolate S130 (AWIW01000000), S134 (AWIS01000000), 090-96 (JFFP01000036) (33) and MAVP-Q (Accession No. MDWT00000000). The map of *recA* region of the five isolates was illustrated using Easyfig (51).

### Core genome SNP determination and phylogenetic analysis

Whole genome phylogenies were constructed with single nucleotide polymorphisms (SNPs) identified from draft genomes using kSNP3 to produce aligned SNPs in FASTA format (52). A maximum likelihood (ML) tree was then built from the FASTA file using raxMLHPC with model GTRGAMMA and the ‐f option, and 100 bootstraps (53). Since there were no differences among the clade II ST631 isolates we used a subset representing geographic and temporal span of isolation.

Minimum spanning tree (MST) analysis was built based on core gene SNPs produced from a cluster analysis. The cluster analysis of ST631 was performed using a custom core genome multi-locus sequence type (cgMLST) analysis using RidomSeqSphere+software v3.2.1 (http://www.ridom.de.seqsphere, Ridom GmbH, Münster, Germany) as previously described (31). Briefly, the software first defines a cgMLST scheme using the target definer tool with default settings using the PacBio generated MAVP-Q genome as the reference. Then, five other *V. parahaemolyticus* genomes (BB22OP, CDC_K4557, FDA_R31, RIMD2210633, and UCM-V493) were used for comparison with the reference genome to establish the core and accessory genome genes. Genes that are repeated in more than one copy in any of the six genomes were removed from the analysis. Subsequently, a task template was created that contains both core and accessory genes. Each individual gene locus from MAVP-Q was assigned allele number 1. Then each ST631 isolate genome assembly was queried against the task template, where any locus that differed from the reference genome or any other queried genome was assigned a new allele number. The cgMLST performed a gene-by-gene analysis of all core genes (excluding accessory genes) and identified SNPs within different alleles to establish genetic distance calculations.

### Configuration and distribution of VPaIs

The VPaI sequence from the PacBio sequenced genomes of MAVP-Q and MAVP-R were identified by comparison with the published RIMD2210633 VPaI-7 (NC_004605 region between VPA1312 – VPA1395) and VPaIxmm (AB455531) (16). Identification of the complete MAVP-Q VPalγ and genomic junctures in chromosome II was done by comparison with the same region of chromosome II in MAVP-R and G149 (which lack an island in this location) using Mauve (54). In a reciprocal manner, the absence of an island in chromosome I in MAVP-Q and G149 was assessed by comparison with chromosome I of MAVP-R. MAVP-Q VPaly (MF066646) and MAVP-R VPaIβ (MF066647) were then extracted as a single contiguous sequence and annotated using Prokka 1.9. Gene content and order of the VPaI elements in MAVP-Q, MAVP-R and RIMD2210633 were then illustrated by Easyfig (51). Roary (55) was then employed to determine homologs among VPaIs based on each island’s annotated sequences with identity set at 50%. Identification of the genome locations of VPaIβ in ST1127 isolate MAVP-M (accession number GCA_001023155) and for VPaFf in AQ4037 (accession number GCA_000182365) (17) was also done using Mauve (54).

To examine the distribution of the VPaIγ in all publicly available draft genomes (https://www.ncbi.nlm.nih.gov/genome/genomes/691, 2016) and genomes from archived regional isolates, whole draft genome sequences were aligned to a 6,118 bp subsequence of the MAVP-Q VPaI with NASP version 1.0.2 (56) (https://pypi.python.org/pypi/nasp/1.0.2, 2017). This subsequence spanned the unique juncture of the four conserved hypothetical proteins (AB831_22090, AB831_22095, AB831_22100, AB831_22105) with the adjacent inserted *tdh* (AB831_22110, c.a. 2549 bp upstream of *ure* cluster)(Supplemental Fig. 1). Percent coverage of the reference sequence was used to determine whether each genome harbored only the four hypothetical proteins, only a *tdh* gene, or the entire module including the fusion of the four genes with *tdh* (Supplemental Fig. 1 and Supplemental Table 3). The sequence type of each genome harboring the fused element characteristic of VPaFγ was then determined using the SRST2 pipeline (48). Where sequencing reads were not available as the input for SRST2, they were simulated from assemblies using an in-house Python script (https://github.com/kpdrees/fasta2reads).

A PCR amplification approach was developed and applied to survey the presence of *tdh* adjacent to the *ure* gene cluster. Primers were designed to conserved sequences of the 3' end of *tdh* (PIHybF8: 5'GCCAACATGGATATAAATAAAAATGA3 ') and the 5' end of *ureG* (tdhUreGrev5: 5'GACAAAGGTATGCTGCCAAAAGTG3') as determined by gene alignments, which when used together produced a 2631 bp amplicon of the insertion juncture when used with MAVP-Q as a template (Supplemental Fig. 4). Amplification was performed on purified DNA with Accustart enzyme mix, with an initial denaturation at 94°C for 3 min., followed by 30 cycles of a denaturation at 94°C for 1 min, annealing at 61°C for 1min, and amplification at 72°C for 2.5 min, with a final elongation at 72°C for 5 min. This amplification was performed in parallel with a diagnostic multiplex PCR amplification of *tdh, trh* and *tlh* using published methods (10, 57) to investigate the co-occurrence of VPalγ with both hemolysin encoding genes in representative isolates of various clinically prevalent sequence types. Amplicons were visualized using a 1.2% agarose gel in TAE buffer (Supplemental Fig. 4).

### Nucleotide sequence accession numbers

The accession number of Pacific Biosciences sequenced genome for MAVP-Q is MDWT00000000, and for MAVP-R is MPPP00000000. The accession number of Illumina sequenced draft genome for G6928 is MPPN00000000, for MA561 is MPPM00000000 and for G149 is MPP000000000. Detailed information about all other ST631 isolate draft genomes were described previously (31) and are listed in Supplemental Table 1. The accessions for the short reads for the remaining sequenced genomes are listed in Supplemental Table 4. The accession number of VPalβ from MAVP-R is MF066647 and the accession number of VPalγ from MAVP-Q is MF066646.

## ACKNOWLEDGEMENTS

We are grateful for clinical isolates and wish to thank specifically: Jana Ferguson and Tracy Stiles of the Massachusetts Department of Public Health, and M. Hickey and C. Schillaci from the Massachusetts Department of Marine Fisheries; J.K. Kanwit of the Maine Department of Marine Resources and A. Robbins from the Maine Department of Health and Human Services; and Laurn Mank from the Connecticut Department of Public Health Laboratory, and K. DeRosia-Banick, Connecticut Department of Agriculture, Bureau of Aquaculture. Assistance with genome sequencing was provided by W. K. Thomas, and technical assistance provided by J. Lemaire, K. Hartman, C. Hallee, M. Malanga, S. Ilyas, J. Hall, J. Sevigny, M. Dillon, K. Flynn, A. Goupil, J. Means, R. Foxall, E. DaSilva, and M.S. Pankey. Partial funding for this work was provided by the USDA National Institute of Food and Agriculture (Hatch projects NH00574, NH00609 [accession number 233555], and NH00625 [accession number 1004199]). Additional funding was provided by the National Oceanic and Atmospheric Administration College Sea Grant program and grants R/CE-137, R/SSS-2, and R/HCE-3. Support was also provided through the National Institutes of Health (1R03AI081102-01), the National Science Foundation (EPSCoR IIA-1330641), and the National Science Foundation (DBI 1229361 NSF MRI). N.G.-E. was funded through the FDA Foods Science and Research Intramural Program. Feng Xu and Cheryl A. Whistler declare a potential conflict of interest in the form of a pending patent application (U.S. patent application 62/128,764). This is Scientific Contribution Number 2722 for the New Hampshire Agricultural Experiment Station.

